# Umbelliprenin isolated from *Ferula sinkiangensis* inhibits tumor growth and migration through the disturbance of Wnt signaling pathway in gastric cancer

**DOI:** 10.1101/455949

**Authors:** Lijing Zhang, Xiaobo Sun, Jianyong Si, Guangzhi Li, Li Cao

## Abstract

**The traditional** herb medicine *Ferula sinkiangensis* K. M. Shen **(*F.* sinkiangensis)** has been used to treat stomach disorders in Xinjiang District for centuries. Umbelliprenin is the effective component isolated from *F. sinkiangensis* which is particularly found in plants of the family Ferula. We previously reported the promising effects of Umbelliprenin against gastric cancer cells, but its **anti-migration** effect **remained** unknown. Here we investigated the **anti-migration** effect and mechanism of Umbelliprenin in human gastric cancer cells. In SRB assay, Umbelliprenin showed cytotoxic activities in **the** gastric cancer cell lines AGS and BGC-823 in a dose-and-time-dependent manner, while it showed **lower** cytotoxic activity in **the** normal gastric epithelium cell **line** GES-1. During transwell, scratch and colony assays, the **migration** of tumor cells was inhibited **by** Umbelliprenin treatment. **The expression levels of the Wnt-associated signaling pathway proteins were analyzed with western blots, and the results showed that** Umbelliprenin decreased the expression levels of **proteins of the Wnt signalling pathway, such as** Wnt-2, β-catenin, GSK-3β, p-GSK-3β, Survivin and c-myc. The translocation of β-catenin to the nucleus was also inhibited **by** Umbelliprenin treatment. In TCF reporter assay, **the transcriptional activity of T-cell factor/lymphoid enhancer factor (TCF/LEF) was** decreased after Umbelliprenin treatment. **The** *in vivo* results suggested that Umbelliprenin **induced little to no** harm in **the** lung, heart and kidney. Overall, these data provided evidence that Umbelliprenin may inhibit the growth, invasion and **migration** of gastric cancer cells **by disturbing the** Wnt signaling pathway.

## 1. Introduction

Gastric cancer is the fourth most common cancer worldwide and **is one of** the most prevalent **cancers among** men in China [Kim et al., 2015; Zheng et al., 2016], which may be **related to their preference for** pickled food and red meat [Zeng et al., 2015]. The clinical outcome of gastrointestinal cancer surgery is limited **because of the** high rates of postoperative complications**, such** as **the** systemic inflammatory response [Climent et al., 2015], which may increase tumor recurrence after surgery [Bohle et al., 2010]. Therefore, **studies have focused** on raising the overall survival rate for gastric cancer [Cho et al., 2016]. However, only trastuzumab and ramucirumab **have been** approved for the treatment of advanced gastric cancer as targeted monoclonal antibodies [Roviello et al., 2016], suggesting that less toxic and more effective therapeutic options are necessary.

Metastasis is the main cause of death in cancer patients [Tang et al., 2014]. The invasion of cancer cells is affected and modulated by many biological molecules and signaling pathways. Growing **evidence indicates that there are abnormalities in the Wnt pathway** in human gastric cancer [Chiurillo, 2015]. In the absence of Wnt signals, β-catenin in the cytosol is continuously degraded. On the contrary, in the presence of Wnt signals, the level of β-catenin increases, **and then,** β-catenin accumulates in the cytosol and subsequently translocates to the nucleus **and**, binds to the T-cell factor (TCF)/lymphoid enhancer factor (LEF). **These** transcription factors regulate the expression of specific downstream target genes such as c-myc and Survivin, which are involved in oncogenesis [Chen et al., 2001; Dong et al., 2015]. Therefore, the **components of the Wnt signalling pathway could be good molecular targets** for gastric cancer therapy.

Natural products have been used as traditional medicines for gastric cancer therapy [Yang et al., 2013]. *Ferula sinkiangensis* K. M. Shen is a traditional folk medicine, which **has** been used for **treating stomach disorders** in Xinjiang District since **the** Tang Dynasty. Umbelliprenin is an effective component **of** *F. sinkiangensis* and exhibits anti-cancer effects in many cancer cells lines [Ziai et al., 2012; Mousavi et al., 2015]. Our previous studies have first explored the growth inhibition effect of Umbelliprenin in gastric cancer cells**: this compound** could induce apoptosis and cell cycle arrest [Zhang et al., 2015]. In the present report, we reported that Umbelliprenin **can inhibit tumor** growth and **migration**, and **that** the underlying anticancer mechanism of Umbelliprenin is associated with **the** Wnt signal pathway.

## 2. Material and methods

### 2.1 Reagents and antibodies

Dulbecco’s Modified Eagle’s Medium (DMEM), Ham’s F12 medium, trypsin, penicillin, streptomycin **and** fetal bovine serum (FBS) were purchased from Gibco (CA, USA). **Sulforhodamine B (SRB)** and **Dimethyl Sulphoxide (DMSO)** were purchased from Sigma-Aldrich (MO, USA). Transwell systems were purchased from Corning (NY, USA). Matrigel basement membrane matrix was bought from BD Biosciences (NJ, USA). M50 Super 8x TOPFlash (12457), M51 Super 8x FOPFlash (TOPFlash mutant) (12457) and pRL-SV40P (27163) were obtain from Addgene. Lipofectamine 2000 was **bought** from Thermo (MA, USA). The dual luciferase reporter assay kit was obtained from KeyGEN Biotech (Jiangsu, China). The nuclear and cytoplasmic protein extraction kit was purchased from Beyotime Biotechnology (Jiangsu, China). Antibodies against Wnt-2, GSK-3β, c-myc, β-catenin, p-GSK-3β, Survivin, Lamin B and β-actin were purchased from Cell Signaling Technology (MA, USA). The cECL Western Blot Kit was obtained from CoWin Biotech (Beijing, China). All the chemical reagents were of the highest grade.

### 2.2 Cell culture and compounds

**The** human gastric carcinoma cell line AGS was cultured in Ham’s F12 medium containing 10% FBS, 100 U/mL penicillin, and 100 μg/mL streptomycin at 37°C with 5% CO2. The human normal gastric epithelial cell line GES-1 **and** human gastric cancer cell line BGC-823 were cultured in DMEM supplemented with 10% FBS, 100 U/mL penicillin and 100 μg/mL streptomycin under the same conditions. Cells were passaged at least three times before being used in experiments. Umbelliprenin was obtained from the seeds of *F. sinkiangensis* as previously described [Zhang et al., 2015]. Umbelliprenin was dissolved as stock solutions in dimethyl sulfoxide **that were** diluted with medium prior to use so **that** the final concentration of DMSO was less than 0.1% (v/v).

### 2.3 Animals

Six-week-old male BALB/c nude mice were obtained from **the** Vital River Laboratories (Beijing, China) and maintained **in** a 12 h light/dark cycle environment (25 ± 2°C) where they **received** and food ad libitum. The protocol for **the** animal **experiments** was approved by the Animal Ethics Committee at the Institute of Medicinal Plant Development, Chinese Academy of Medical Sciences.

### 2.4 Cell viability assay

**The** Sulforhodamine B (SRB) assay was used to determine cell viability. Cells were seeded in 96-well plates in triplicate and cultured for 24 h at 37°C. Then, the cells were treated with Umbelliprenin in various concentrations (**0**, 3.125, 6.25, 12.5, 25, 50 μM). After **the** 24 h treatment, 50 μL **of** cold TCA was added to fix **the** cells for 1 h at 4°C. The plates were washed with water and air-dried. **One-hundred microlitres** of 0.4% (w/v) SRB was added and **cells were** stained for 15 min. After washing 4 times with acetic acid, 100 μL **of** Tris-base was added for 10 min, **while shaking**. The absorbance was measured at 540 nm using a Microplate Reader (Bio Tek, USA). % Cytotoxicity= (Control – Experimental) / Control* 100%.

### 2.5 Colony Formation Assay

**The** colony formation **assay** was used to evaluate the anchorage-independent growth of gastric cancer cells. Cells were seeded in 6-well plates and treated with Umbelliprenin (0, 6, 12, 24 μM for AGS **cells** and 0, 12.5, 25, 50 μM for BGC-823 **cells**) for 10 days. The tumor colonies were observed counted **using a** microscope. Then, the colonies were stained with crystal violet (**1 mg/mL**), and colonies larger than 200 μm were counted.

### 2.6 Wound healing Assay

Cell motility was detected using **the** wound healing assay. Cells were cultured in **a** 24-well plate to nearly 90% confluence. The monolayers were then carefully scratched using a sterile 200-μL pipette tip with a constant width. The cells were washed with PBS and treated with Umbelliprenin (IC_50_ values **were used** in both AGS and BGC-823 **cells**). The cells were photographed at 0, 24, 36 and 48 h after treatment to observe the distances **that the cells had migrated**. The cell motility was calculated as **follows**: Cell motility = (distance at 0 h - distance at 24, 36 or 48 h) / distance at 0 h* 100%.

### 2.7 Transwell-migration / invasion assays

Cell migration was analyzed in a 24-well transwell plate with 8-μm pore size polyvinylidene filter membrane. Cells were seeded into the upper chamber (1×10^5^) in serum-free Ham’s F12 **medium** and treated with Umbelliprenin (0, 6, 12, 24 μM for AGS **cells** and 0, 12.5, 25, 50 μM for BGC-823 **cells**). Ham’s F12 **medium** with 10% serum was added to the lower chamber. After incubation for 24 h, **the** cells on the upper side of the filter were removed with cotton swabs, **and** the **filters** were fixed with 4% formaldehyde for 10 min at room temperature, **and** stained with crystal violet for 15 min, the number of cells in five random fields of each triplicate filter **was counted** under the light microscope. The cell invasion assay was conducted under similar procedure, except that 80 μL of Matrigel was used to coat **the** upper chamber for 12 h before the cells were seeded.

### 2.8 β-Catenin / TCF Transcription Reporter Assay

**The** TCF-reporter plasmids (TOPFLASH and the negative control FOPFLASH) and Renilla luciferase (pRL-SV40P) plasmid were obtained from Addgene. Cells were seeded in 6-well plates and the reporter plasmids **containing** TCF binding sites (TOPFLASH, 500 ng/well) or mutant, inactive TCF binding sites (FOPFLASH, 500 ng/well) **were** transiently transfected into **the** cells for 6 hours using Lipofectamine 2000. The cells were co-transfected with **the** Renilla luciferase (pRL-SV40P, 5 ng/well) plasmid to normalize **the** transfection efficiency **with** the internal control. Then, the medium was renewed and different concentrations of Umbelliprenin were added. After **the** 24 h treatment, TCF-mediated gene transcription was expressed by the ratio of TOPFLASH: FOPFLASH luciferase activity, and each **value** was normalized to the Renilla luciferase activity.

### 2.9 Protein isolation and western blot

Fractionated nuclear and cytosolic proteins were obtained according to the manufacturer’s instructions. Briefly, cells were treated with Umbelliprenin (0, 6, 12, 24 μM for AGS **cells** and 0, 12.5, 25, 50 μM for BGC-823 **cells**) for 24 h, harvested and washed twice with ice-cold PBS. 500 μ**L of** cytoplasmic extract agent A and B were added and **the suspension was** centrifuged at 12,000 rpm for 5 min. The supernatant **contained** cytosolic **proteins**. The nuclear residue was mixed with 200 **μL** of **the** nuclear extract agent. The mixture was vortexed for 30 min and then centrifuged at 14,000 rpm for 10 min at 4°C. The supernatant **contained** nuclear **proteins**.

To obtain whole-cell lysates, cells were lysed in lysis buffer after **treatment** with Umbelliprenin (0, 6, 12, 24 μM for AGS **cells** and 0, 12.5, 25, 50 μM for BGC-823 **cells**) for 24 h and **the** protein concentrations were determined by the BCA method. Protein samples were separated by SDS-PAGE and transferred onto PVDF membranes. After blocking for 1 h in **the** 5% non-fat milk solution, the membranes were incubated with **the** primary antibody overnight at 4°C. Then the primary antibody was washed with TBST **for** 3 times and incubated with secondary antibody for 1 h at room temperature. Protein bands were detected using ECL and the levels of β-actin were used to ensure equal loading of proteins.

### 2.10 Immunofluorescence staining to detect the nuclear translocation of β-catenin

Cells were cultured in plates and treated with Umbelliprenin (0, 6, 12, 24 μM for AGS **cells** and 0, 12.5, 25, 50 μM for BGC-823 **cells**) for 24 h. The treated cells were fixed with 4% paraformaldehyde for 10 min and washed **twice with PBS**. These cells were blocked with 5% bovine serum albumin in PBS for 60 min, and then incubated with **the** primary antibody (diluted 1:200 in PBS containing 3% BSA) overnight at 4°C. After washing twice with PBS, the cells were co-incubated with **the** IgG PE-conjugated secondary antibody (diluted 1:400 in PBS containing 3% BSA) and DAPI for 1 h at room temperature. The cells were examined with **the** Image Xpress system (Molecular Devices, USA).

### 2.11 Safety of Umbelliprenin in tumor xenograft models

Animal **experiments were** conducted according to **the** previous description [Zhang et al., 2015]. Briefly, mice were inoculated subcutaneously with **the** human gastric cancer cells BGC-823 (1.0 × 10^6^) on the right flank. The mice were randomized to six groups and **their conditions were observed** every day. **Umbelliprenin were diluted in 200 μL 0.9% NaCl solution and administered to each mouse at a dose of 10 mg/kg or 20 mg/kg twice daily for 12 days.** Mice were then euthanized and **the** lungs, livers, hearts and kidneys were fixed and preserved in formalin for hematoxylin-eosin staining.

### 2.12 Statistical analysis

All data were analyzed by **the** IBM SPSS statistics 19 **software**. All tests were conducted at least three times. Statistical significance was defined as *p < 0.05 and **p < 0.01. The results were expressed as mean ± SD which represent three independent tests.

## 3. Results

### 3.1 Umbelliprenin reduced the viability in AGS and BGC-823 human gastric cancer cells, but not in GES-1 cells

The chemical structure of Umbelliprenin isolated from the seeds of *F. sinkiangensis* **is shown** in **Figure 1A**. **Because** Umbelliprenin **appeared to be** most effective in gastric cancer **cells** among other common cancer cells, we studied **its** anti-proliferative effects **in the** two human gastric cancer cell lines: AGS **and** BGC-823 **as well as in the** human gastric epithelial cell line GES-1. The IC_50_ values for the three cancer cell lines varied. The results showed that Umbelliprenin **inhibited** the growth in both **the** AGS and BGC-823 gastric cancer cell lines with an IC_50_ of 11.74 μM and 24.62 μM, **respectively,** the IC_50_ for AGS was lower **than that for the BGC-823 cells** (**Figure 1C** and **Figure 1D**). In addition, Umbelliprenin was less cytotoxic **in** GES-1 cells (IC_50_: 97.55 μM) compared with AGS and BGC-823 cells (**Table 1**). Based on **the** IC_50_ values, we chose 12 μM **and** 24 μM Umbelliprenin**, as well as their multiples or fractions, as the concentrations for treating** AGS and BGC-823 cells, **respectively**. Furthermore, as the colony formation ability is closely related to tumorigenesis *in vivo*, we detected the impact of Umbelliprenin on the growth of gastric cancer cells. The results showed that Umbelliprenin **significantly** inhibited the colony forming ability of AGS **cells** in a dose-dependent manner, while **it was** less effective in BGC-823 cells (**Figure 1B**). Together, these data indicate that Umbelliprenin may decrease the proliferation and tumor forming ability **in AGS and BGC-823 gastric cancer cells**.

**Figure 1:**
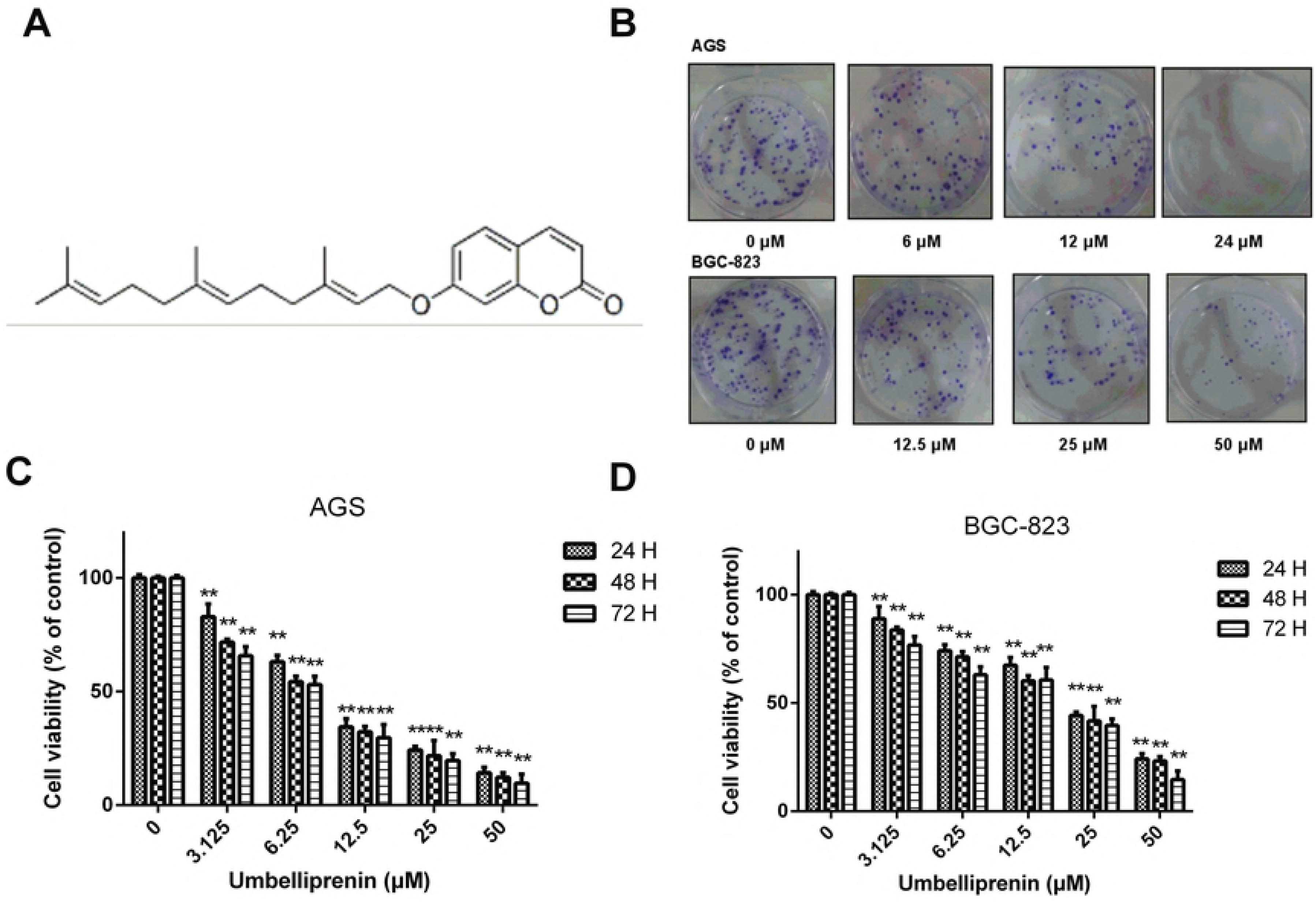
Chemical structure and inhibition effects on cell viability of Umbelliprenin. **A.** Chemical **structure** of Umbelliprenin isolated from the **seeds** of *Ferula sinkiangensis*. **B.** Colony formation assay. AGS or BGC-823 cells were **grown** for 10 days after incubation with different concentrations of Umbelliprenin. **C.** The effects of Umbelliprenin on the **viability of AGS human gastric cancer cells**. **D.** The effects of Umbelliprenin on the viabilities of human gastric cancer cells BGC-823. AGS and BGC-823 **cells** were exposed to various concentrations of Umbelliprenin (0, 3.125, 6.25, 12.5, 25, 50 μM) for 24 h, 48 h or 72 h, followed by the Sulforhodamine B (SRB) assay. The data represent the mean value of three independent experiments and are expressed as **the mean** ± SD. **p < 0.01, *p < 0.05 were considered statistically **significant**.

**Table 1.** Cytotoxicity of Umbelliprenin isolated from the seeds of *Ferula sinkiangensis* in three cell lines.

| Compounds | IC<sub>50</sub>(μM)<sup>a</sup> |  |  |
| --- | --- | --- | --- |
|  | AGS | BGC-823 | GES-1 |
| Umbelliprenin | 11.74±1.33 | 24.62±2.45 | 97.55±3.81 |

<sup>a</sup> IC<sub>50</sub> is the concentration of compound causing 50% growth inhibition for each cell line after 24 h treatments. The results represent the mean values of three independent tests.

### 3.2 The effects of Umbelliprenin on the invasion and migration in AGS and BGC-823 cells

Umbelliprenin was effective in reducing cellular migration in gastric cancer cells. **We performed a wound healing assay, and** images representing **the migration capability of the cells were taken at different time points,** at the same **site and magnification**. **Forty-eight hours after scratching**, **in the presence of Umbelliprenin at its IC_50_ value,** the wound was healed approximately **by** 77.4% and 64.2% for AGS and BGC-823 **cells** compared with **the** control group, respectively **(Figure 2C)**. The migration assay using the transwell-migration system also showed that Umbelliprenin **effectively inhibited** the migration of AGS and BGC-823 **cells (Figure 2A** and **Figure 2B)**. Furthermore, the inhibitory effect of Umbelliprenin on the invasion of gastric cancer cells was examined **in a transwell assay using** Matrigel-coated filters. Compared with **the** control group, **fewer** AGS and BGC-823 cells **penetrated** the filters after Umbelliprenin treatment**. Additionally,** Umbelliprenin decreased the invasive potential of cells in a dose-dependent manner **(Figure 2A** and **Figure 2B)**. **Therefore, both the** migration and invasion of gastric cancer cells were significantly suppressed after Umbelliprenin treatment.

**Figure 2:**
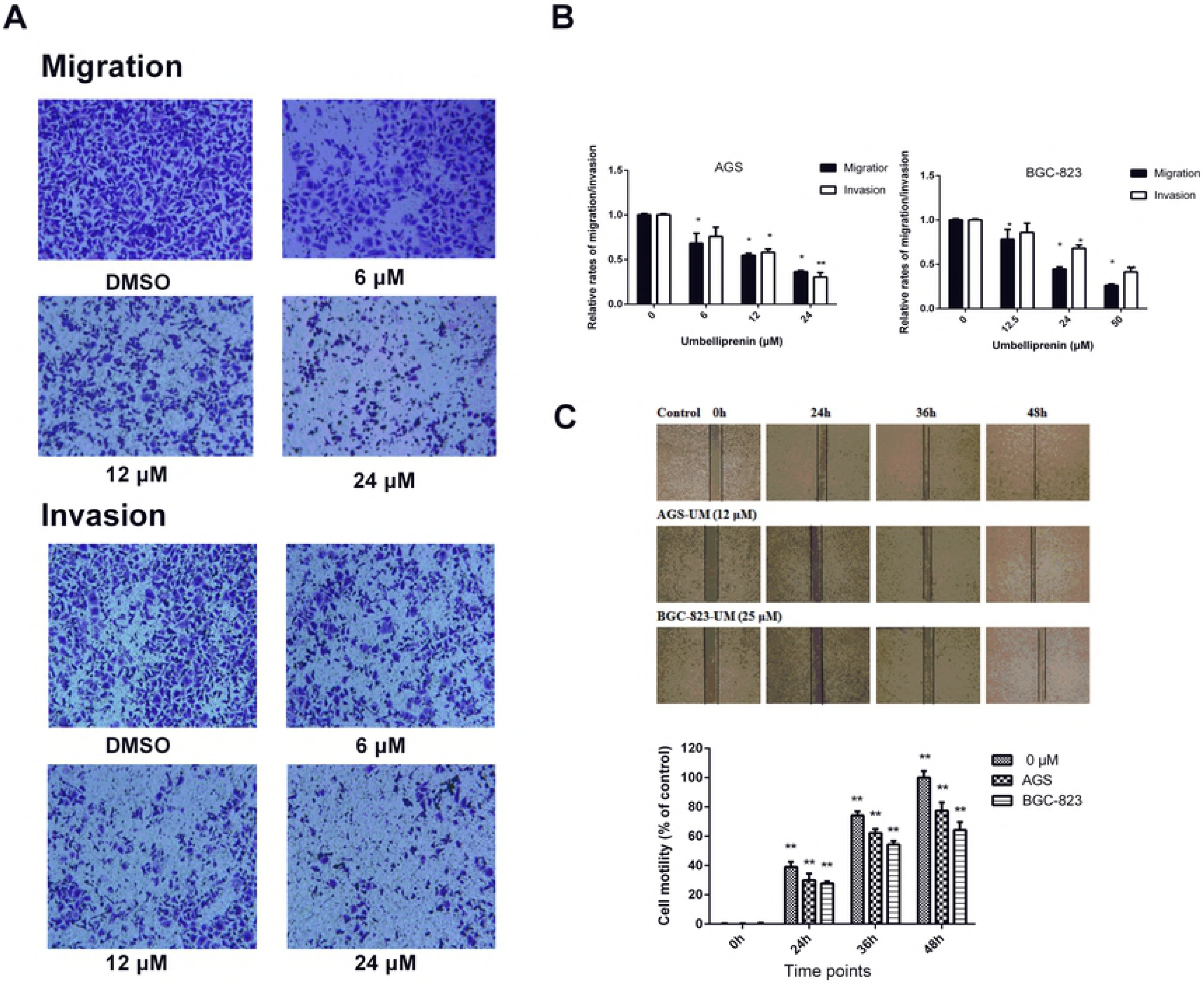
Umbelliprenin inhibits the migration and invasion of gastric cancer cell lines. **A.** The migration or invasion of cells was measured by transwell or matrigel-coated filter transwell assays, **respectively**. Cells were incubated with Umbelliprenin for 24 h, and the migrated or invaded cells were fixed and stained with crystal violet. **B.** The migration and invasion assays data are expressed as the means ± SD and represent the mean value of three independent **experiments**. **p < 0.01, *p < 0.05. **C.** The migration of AGS and BGC-823 **cells was examined in a** wound healing assay. Cells were scratched and treated with Umbelliprenin. Images were **obtained using** a microscope at 0 h, 24 h, 36 h and 48 h. Data are presented as the **mean** ± SD and represent three independent **experiments**. **p < 0.01, *p < 0.05 were considered statistically significance.

### 3.3 Umbelliprenin inhibited the transcriptional activity of β-catenin/TCF in the human gastric cancer cell lines AGS and BGC-823

To explore **whether Umbelliprenin plays a role on the dysregulation of the** Wnt/β-catenin pathway in gastric cancer cells, we examined the **transcriptional** activity of β-catenin/TCF after Umbelliprenin treatment. **For the purpose, we transfected the** TOPFLASH or FOPFLASH **plasmids** into AGS and BGC-823 gastric cancer cells. The transfection efficiency was normalized **with Renilla** luciferase. As shown in **Figure 3A** and **Figure 3B**, **after 24 h of treatment** Umbelliprenin reduced the **TCF**-dependent luciferase **activity** (TOPflash) in a dose-dependent manner in both gastric cancer cell lines, while it **did not change the activity of the FOPflash control plasmid**. In AGS cells, the **transcriptional activity of TCF was** 62.93%, 31.54%, and 18.19% **relative to** the control group at concentrations of 6, 12 and 24 μM of Umbelliprenin, respectively (P<0.05). In BGC-823 cells, the **transcriptional activity of TCF was** 52.91%, 43.50%, and 21.15% **relative to** the control group at concentrations of 12.5, 25 and 50 μM of Umbelliprenin, respectively (P<0.05). This indicates that the transcriptional activity of β-catenin/Tcf could be inhibited by Umbelliprenin. **Thus, Umbelliprenin suppresses** the Wnt/β-catenin signaling pathway in the two gastric cancer cell lines.

**Figure 3:**
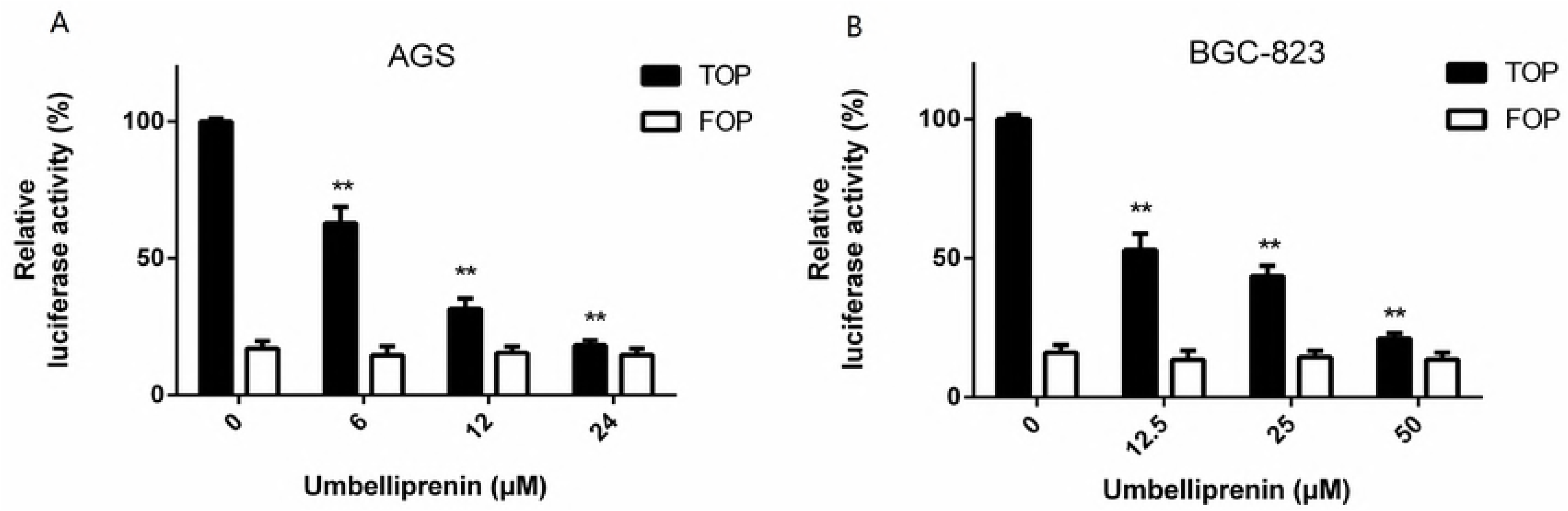
Umbelliprenin treatment decreases the activity of the TCF reporter in gastric cancer cells. **A.** and **B.** AGS and BGC-823 **cells** were transfected with **the** TOPFLASH or FOPFLASH **plasmid** together with **the Renilla plasmid** as control. Cells were then treated with Umbelliprenin (0, 6, 12, 24 μM for AGS **cells** or 0, 12.5, 25, 50 μM for BGC-823 **cells**) for 24 hours. **The luciferase activity was measured and the results were normalized.** The data represent the mean value of three individual experiments. *p < 0.05 and **p < 0.01 were considered statistically **significant.**

### 3.4 Inhibition of nuclear translocation of β-catenin by Umbelliprenin

To determine whether Umbelliprenin suppressed β-catenin mediated transcription **by interfering with the nuclear translocation of β-catenin**, we investigated the β-catenin subcellular localization in Umbelliprenin -treated AGS and BGC-823 cells by immunofluorescence staining. As shown in **Figure 4A**, β-catenin preferentially accumulated in the nucleus in **the** control group. **By** contrast, after treatment with Umbelliprenin for 24 h, the localization of β-catenin decreased **in the nucleus but increased in** the cytoplasm and at the plasma membrane. This result was confirmed by Western blot analysis in which β-catenin was increased in the cytoplasmic **protein** fraction **and** decreased in the nuclear **protein** fraction **of the** treated groups **(Figure 4B)**. **The** relative expressions of cytoplasmic and nuclear proteins were shown in **Figure 4C**. Therefore, Umbelliprenin may mediate its effect by inhibiting the translocation of β-catenin to the nucleus.

**Figure 4:**
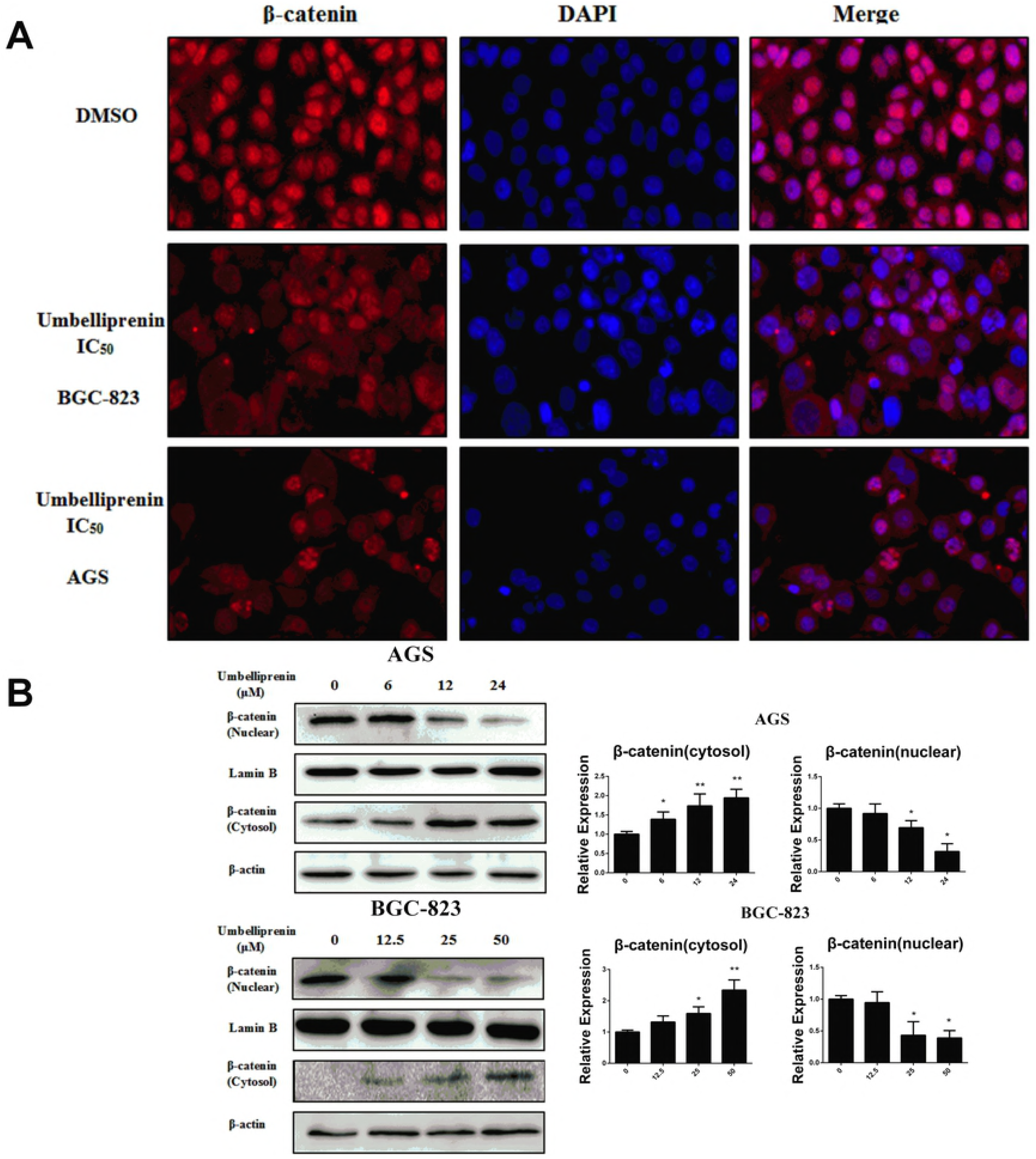
Umbelliprenin inhibits the nuclear translocation of β-catenin in gastric cancer cells. **A.** The translocation of β-catenin was examined by immunostaining. Cells were treated with UM (0, 6, 12, 24 μM for AGS **cells** or 0, 12.5, 25, 50 μM for BGC-823 **cells**) for 24 h, stained with **the** anti-β-catenin antibody and **analysed** using **the** Image Xpress Micro imaging system (MD, USA). **B.** Protein expressions were detected by western blot. After treatment with Umbelliprenin (0, 6, 12, 24 μM for AGS **cells** or 0,12.5, 25, 50 μM for BGC-823 **cells**) for 24 h. Cellular fractionation was carried out to determine the cellular localization of β-catenin. Lamin B and β-actin were used as **controls for** nuclear fraction and cytoplasmic fraction, **respectively**. **C. The** relative expression of cytoplasmic and nuclear proteins was analyzed. **The** results obtained from a **representative** experiment are shown (n=3). Statistical significance was **p < 0.01.

### 3.5 Umbelliprenin inhibits Wnt signaling by decreasing the phosphorylation of GSK-3β and reducing the downstream effectors of Wnt signaling

Wnt signaling substantially impacts gastric tumorigenesis and prognosis [Bohle et al., 2010]. **To** further determine the mechanisms by which Umbelliprenin inhibits cellular proliferation, migration and invasion, we studied the Wnt signaling. The expression levels of Wnt signaling-associated proteins in both AGS and BGC-823 cells were measured by western blot. Wnt-2, β-catenin, and GSK-3β, potential modulators of Wnt signaling, as well as the downstream targets of Wnt signaling Survivin and c-myc, were significantly reduced **in Umbelliprenin treated cells** compared to the control group (**Figure 5A** and **Figure 5C)**. **The** relative expressions were shown in **Figure 5B** and **Figure 5D**.

**Figure 5:**
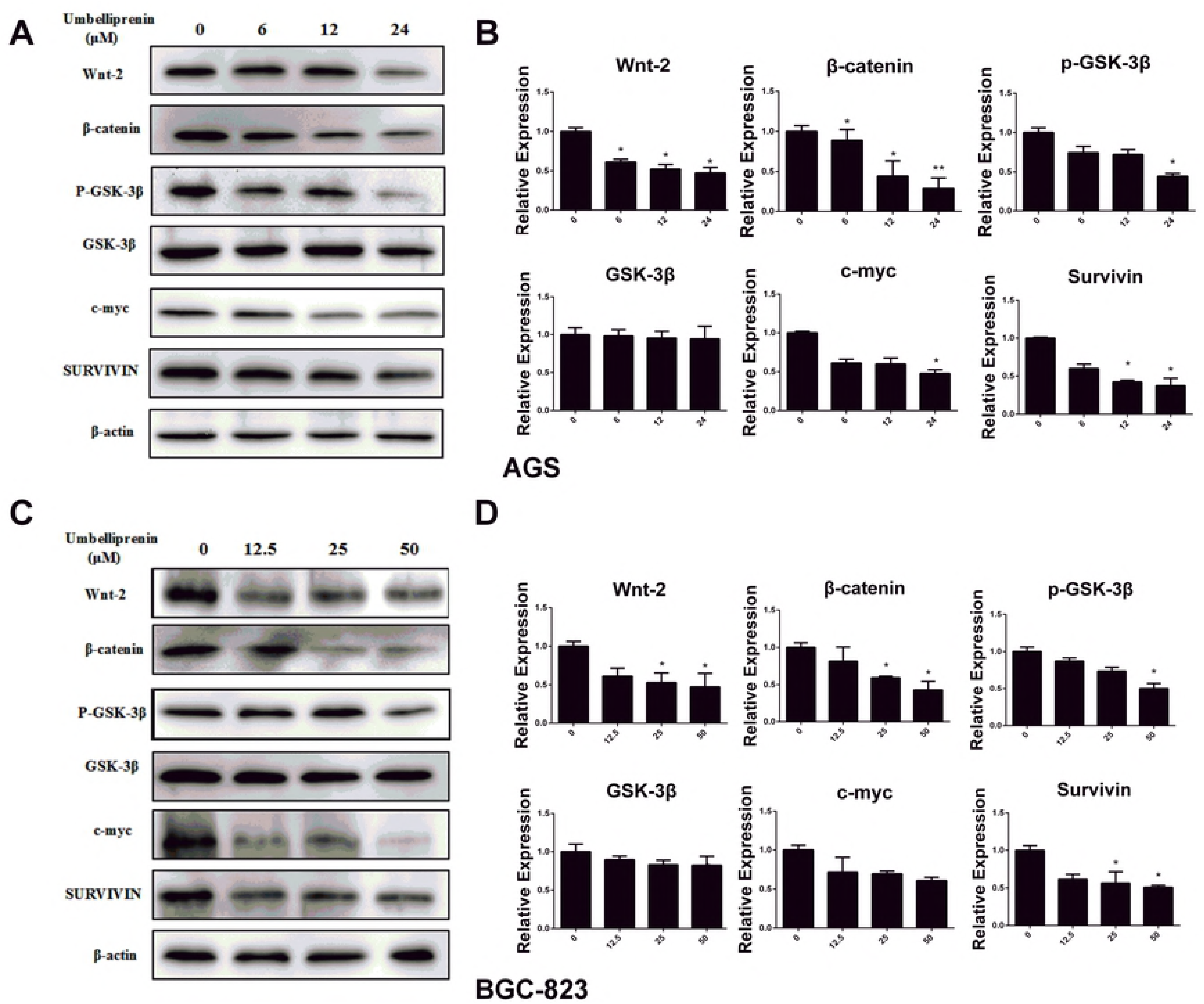
Umbelliprenin downregulates the Wnt signal pathway, and Survivin and c-myc protein expression levels. **The expression of the regulatory proteins of the** Wnt signal pathway, Survivin and c-myc protein were determined by western blot. **A.** and **C.** AGS or BGC-823 cells were treated with Umbelliprenin (0, 6, 12, 24 μM for AGS **cells** or 0, 12.5, 25, 50 μM for BGC-823 **cells**) for 24 h. β-actin was used to confirm equal protein loading. **B.** and **D. The** relative expression levels of proteins are **shown.** All tests were **performed** in triplicate. *p < 0.05 and ** p<0.01 were considered statistically significant.

### 3.6 The safety of Umbelliprenin in BGC-823 tumor xenograft models

Previous **studies** showed that Umbelliprenin could effectively inhibit the growth of tumor *in vivo*. Thus, we evaluated the safety of Umbelliprenin in BGC-823 tumor xenograft models. All mice tolerated the treatment procedure well **and did not show** toxic symptoms or signs. **The analysis of biochemical markers for the liver, as ALT and AST, showed no significant change between the** different groups **(Table 2)**. In addition, no histological abnormality was shown in **the** lungs, liver, heart, and kidneys of mice between **the** Umbelliprenin treatment groups and control group at the end of **the** treatment (**Figure 6**). Together, these data suggest that Umbelliprenin effectively inhibits tumor growth, **and does not cause** obvious drug-induced adverse effects.

**Table 2.** Serum analysis for liver function (mean ± SD, n = 8) Blood samples were collected and centrifuged at 12,000 rpm for 15 min to obtain the serum. Liver function was evaluated based on the serum levels of aspartate aminotransferase (AST) and alanine aminotransferase (ALT). All biochemical parameters were evaluated by an automated biochemical analyzer (Beckman).

| Group | AST(U/L) | ALT(U/L) |
| --- | --- | --- |
| Control | 202.6±27.9 | 31.8±2.3 |
| Umbelliprenin (10mg/kg) | 190.4±33.8 | 30.3±4.1 |
| Umbelliprenin (20mg/kg) | 172.9±17.1* | 32.1±3.1 |

\*\*p < 0.01, \*p < 0.05 were considered statistically significance compared with control group. **And the biochemical markers ALT and AST were shown no significance between different groups.**

**Figure 6:**
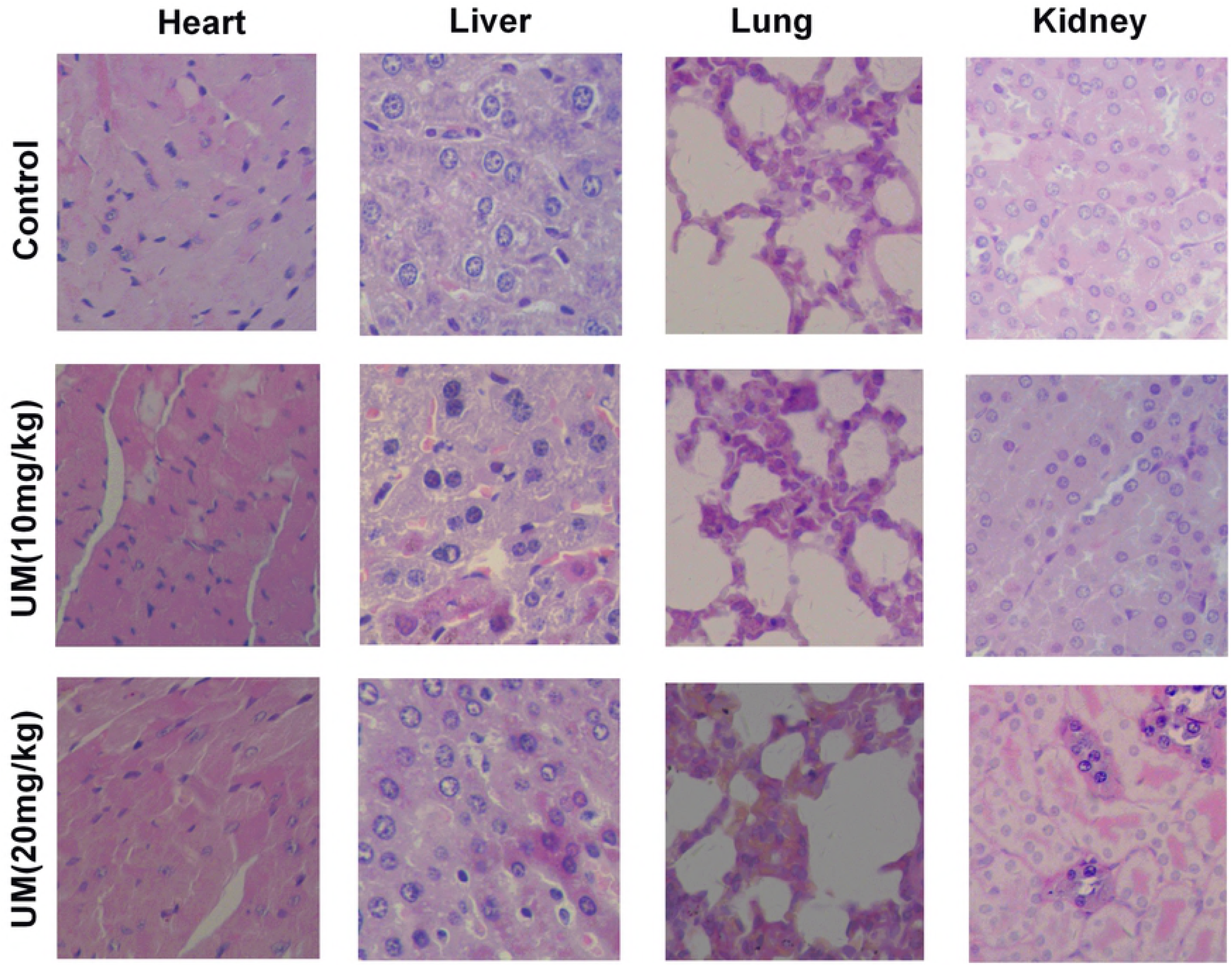
Low toxicity of Umbelliprenin *in vivo*. The results showed representative **hematoxylin-eosin** staining **to investigate the** potential toxicity of Umbelliprenin in lung, heart, kidney and liver of Umbelliprenin -treated and control mice on day 12 **(n = 8).**

## 4. Discussion

Natural products with potential for gastric cancer treatment have **received much** attention. Umbelliprenin has been reported to possess **the** promising effect of inducing apoptosis **in** a variety of cancer cell types such as leukemia, breast cancer and bladder carcinoma [Mousavi et al., 2015; Gholami et al., 2013; Haghighi et al., 2014] and **exhibits** biological effects in multiple signal transduction pathways. For example, **a** study showed that Umbelliprenin could activate both **the** intrinsic and extrinsic pathways of apoptosis in **the** Jurkat T-CLL cell line, which accounted for the inhibition of cellular proliferation and apoptosis induction [Gholami et al., 2013]. **Therefore,** further studies on the **anti-gastric cancer effects of Umbelliprenin** are necessary. Our previous results showed that Umbelliprenin **among seven compounds isolated from the seeds of *F. sinkiangensis*, exhibits the strongest antitumor effect *in vivo* and *in vitro*.** Based on these results, we performed further research on Umbelliprenin. In the present study, we demonstrated the **anti-metastatic and anti-proliferative** effects of Umbelliprenin on **the** AGS and BGC-823 gastric cancer cell lines *in vitro* and its safety *in vivo*. These results indicated that Umbelliprenin might be an effective inhibitor of tumor **migration**, **and were obtained using** wound healing, transwell migration and invasion assays. We also demonstrated that the **mechanism** of action of Umbelliprenin include the inhibition of **the** Wnt signaling pathways **and** the decrease of c-myc and Survivin.

The activation of **the** Wnt/β-catenin signaling is found **approximately** 30% to 50% of gastric cancer tissues and in many gastric cancer cell lines [Ooi et al., 2009]. β-catenin is a multifunctional protein that was found as **an** E-cadherin-binding protein involved in the regulation of **cell-to-cell** adhesion and works as a transcriptional regulator in the Wnt signaling pathway [Kikuchi et al., 2011]. It accumulates in the cytoplasm and translocates to the nucleus, **where it** behaves as transcriptional coregulatory **factor** by interacting with **the** TCF/LEF complex and activates target oncogenes, **such as** c-myc and Survivin [Eastman et al., 1999; Zhang et al., 2014]. Our results suggest that Umbelliprenin down-regulates Wnt, resulting in **the** decrease **of** phosphorylated GSK-3β and reduction of Wnt downstream effectors, **such as** β-catenin, Survivin, c-myc, MMP2 and MMP9. Furthermore, Umbelliprenin treatment caused the inhibition of **the** nuclear translocation of β-catenin and the reduction of **the activity of the TCF-reporter**. Although results suggest that Umbelliprenin **interferes with the** Wnt/beta-catenin signaling and **therefore inhibits the activity of the TCF reporter, reduction of the** reporter activity might be partially caused **the Umbelliprenin-induced** migration and invasion. Our results are consistent with **those** previous studies showing that decreased Wnt expression is associated with decreased phosphorylation of GSK-3β; decreased **expression of** β-catenin, c-myc and Survivin; **lower activity of the TCF reporter; and reduced** migration and invasion [Luo et al., 1999].

Taken together, **for the first time,** we showed strong evidence that Umbelliprenin isolated from the **seeds** of *F. sinkiangensis* can inhibit **cell** growth, migration and invasion, at least in part, through the Wnt signaling pathway. Other reports have shown that the Wnt antagonists inhibit the growth of HCC cells *in vitro* and *in vivo* **through inhibition of the formation of the** TCF-4/β-catenin complex and its transcriptional activity **and** downregulation of **the** β-catenin/TCF-4 target genes c-myc and Survivin [Wei et al., 2010]. **By** contrast, we found that Umbelliprenin inhibits the Wnt pathway by decreasing nuclear translocation of β-catenin rather than **disrupting the** TCF-4/β-catenin complex. Therefore, **treatments that** combine Umbelliprenin with **an** antagonist of TCF-4/β-catenin may lead to promising effects on suppressing the activation of **the** Wnt signaling pathway.

In conclusion, our data indicate that Umbelliprenin **has anti-migration** effects in AGS and BGC-823 gastric cancer cells, **targets** the Wnt signaling pathway, and **exhibits** good safety during the treatment in **the** BGC-823 xenograft *in vivo* **model**: **indeed,** no **abnormalities in regard to** body weight, daily diet, liver function and histological characteristics **of** lung, spleen, heart, kidney and liver tissue were observed. Therefore, Umbelliprenin could be a promising approach for gastric cancer treatment; however, further investigations are necessary.

## Acknowledgments

We would like to thank Addgene for providing us **with the** M50 Super 8x TOPFlash (12457), M51 Super 8x FOPFlash (TOPFlash mutant) (12457) and pRL-SV40P (27163) **plasmids**. This work **was supported** by the National Natural Science Foundation of China (No. 81460661 and No. 81460586), the Ministry of Science and Technology of the People's Republic of China, and **the** Major Scientific and Technological Special Project for “Significant New Drugs Formulation” (Grant No. 2012ZX09501001-004 and 2012ZX09301-002-001-026). This work was also supported by **the** Beijing Key Laboratory of Innovative Drug Discovery of Traditional Chinese Medicine and Translational Medicine, Institute of Medicinal Plant Development, Peking Union Medical College, Chinese Academy of Medical Sciences.

